# Emergence of synchronised growth oscillations in filamentous fungi

**DOI:** 10.1101/2023.12.22.573137

**Authors:** Praneet Prakash, Xue Jiang, Luke Richards, Zoe Schofield, Patrick Schafer, Marco Polin, Orkun S. Soyer, Munehiro Asally

## Abstract

Soil fungi are important decomposers of organic matter and play crucial roles in the biogeochemical cycles in the soil. Many species of fungi grow in the form of branched networks. While there have been investigations on the growth and architecture of the fungal networks, their growth dynamics in space and time is still not fully understood. In this study, we monitor the growth dynamics of the plant-promoting filamentous fungus *Serendipita indica* for several days in a controlled environment within a microfluidic channel. We find that this species displays synchronized growth oscillations with the onset of sporulation and at a period of 3 hours. Quantifying this experimental synchronisation of oscillatory dynamics, we show that the synchronisation can be captured by the nearest neighbour Kuramoto model. Our analysis suggested the existence of millimetre-scale cell-cell communication across the fungi network. The microfluidic setup presented in this work may aid the future characterization of the molecular mechanisms of the cell-cell communication, which could in turn be exploited in order to control fungi growth and reproductive sporulation in soil and plant health management.

## Introduction

Filamentous fungi and some amoeboids have evolved a lifestyle where a single colony incessantly grows and spreads in the form of a network to find nutrient resources [1]. The amoeboid *Physarum polycephalum* is among the best studied organisms for this locomotive network lifestyle. Their locomotion is driven by synchronised peristaltic contraction [2], and the ensuing complex exploratory behaviour enables them to solve spatial optimisation problems in the search for food locations [3–5]. Filamentous growth gives rise to branched networks that can offer a much larger surface area which is desirable for an efficient exchange of nutrients or molecules.

In addition to their spatial coordination, microbes can exhibit coordination in time. For example, *Physarum* networks are known to coordinate their movement over long distances [6], switch between behavioural states and are capable of encoding memory like neurons [7]. Just like *Physarum*, filamentous fungi also form branched growth patterns. But, instead of a peristaltic spread, the individual filaments are rigid and either grow over a surface or bore through the medium. Filamentous fungi can also communicate over long distances [8,9], regulate their growth and can make new connections or fuse branches to re-organise their network architecture [10]. Fungi can exhibit an extremely large-scale locomotive network lifestyle, with largest known fungal network spanning as much as a few square kilometres [11,12].

So far, the studies on fungal networks have largely focused on understanding the mechanisms behind the growth of individual fungal filaments, known as hyphae. The growth of hyphae is linked with internal hydrostatic pressure (turgor) changes, along with hyphae tip ‘softening’ and ‘hardening’ [13,14]. Individual hyphal tips are known to grow periodically, and synchronisation of this periodic growth plays an important role in the fusion of hyphal filaments, and consequently the organisation of network architecture [15]. On the scale of hyphal networks, branching patterns, and the dynamics of nutrient supply within the network have been explored [16–18]. These studies have revealed oscillatory spatial domains of nutrient acquisition within the network [8]. However, the growth dynamics of the global hyphal networks is still largely unexplored. Specifically, it is unclear if fungal growth is determined locally or represents an orchestrated activity of the entire network.

*Serendipita indica* (*Piriformospora indica*) is a fungal root endophyte [22–24]. In addition to a saprophytic lifestyle on dead organic matter, *S. indica* forms symbiotic relationships with bacteria and plants [19,20]. Their relationship with plants encompasses nutrient exchange and protection of plants against biotic and abiotic environmental stress [21]. *S. indica* can colonise a broad range of plants for its reproduction [22]. Hence, it is a useful and relevant model system to investigate the fungal network growth dynamics.

In this work, we monitored the long-term growth of *S. indica* hyphal networks within a microfluidic device, from spore germination until the formation of new spores. The whole process lasted several days, during which we maintain a constant temperature and nutrient media supply in the device. During the first few days, hyphal networks spread uniformly. However, later during the reproductive phase, as spores began to form, the culture exhibited oscillatory growth pattern that synchronised across the network. We explain this synchronisation behaviour using the Kuramoto model, which is a general model for synchronisation of coupled oscillators [23]. The fitting of the model parameters to experimental observations on oscillation and synchronisation dynamics revealed that that the observed synchronisation is best explained with the assumption of a local coupling within an area of ∼1 *mm*^2^. These findings show that a fungi network can achieve synchronous growth across 10s of mm^2^, through couplings that form within the 1mm^2^ range. Future studies can explore the physical and chemical basis of such coupling and the functional role of synchronisation.

## Materials and Methods

### Model organism and spore harvesting

*S. indica* was axenically cultivated on agar plates infused with glucose-based minimum essential growth media at 30°C [19]. Spores were inoculated on an agar plate (1.5% agar, 0.2 mM MgSO_4_, 5 μM CaCl_2_, 10 μM FeSO_4_, 7.5 μM EDTA, 0.5 μM thiamine-HCl and 100 mM glucose in 1x MEM base media). The agar block inoculated with spores in a typical 30-day-old fungal colony was plucked out and vigorously shaken to detach them from hyphae filaments with 0.02% Tween20-deionized water in a 50 mL falcon tube. The harvested spores were then filtered using nylon filters with a pore size of ∼20 μm.

### Loading *S. indica* spores in a microfluidic device

Fungal growth dynamics experiments were carried out in a custom-designed microfluidic device with a large central reservoir of area ∼20 mm^2^ and a depth of 14 μm i.e., nearly five times the diameter of a hyphal tip (Figure 1(b)). Twin inlets and outlets maintain continuous supply of nutrient media and the two pillars in the centre prevent the reservoir from collapsing. The growth medium was composed of 0.2 mM MgSO_4_, 5 μM CaCl_2_, 10 μM FeSO_4_, 7.5 μM EDTA, 0.5 μM thiamine-HCl and 100 mM glucose in 1x MEM base media. To initiate the experiments, harvested spores were flushed into the microfluidic device maintained at 30°C. They were then supplied with growth media at a flow rate of 100 nl/min. The continuous nutrient supply allowed us to record movies over multiple days.

**Figure 1:**
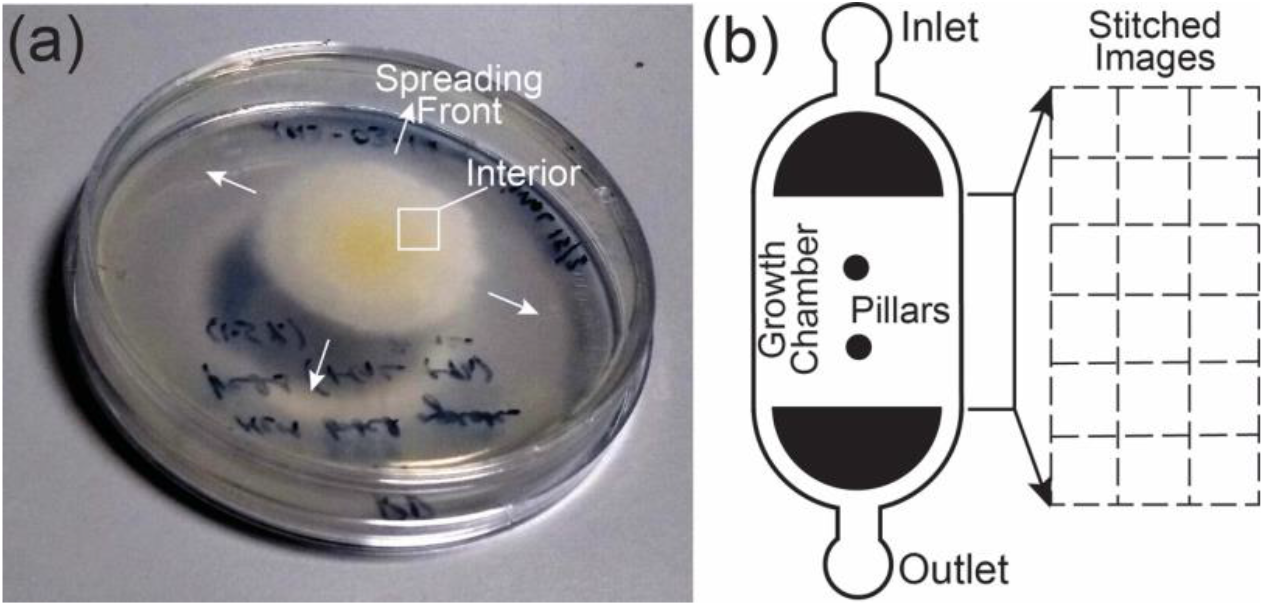
Growing filamentous fungi. (a) *S. indica* colony cultivated on a 55 mm diameter Petri dish for a period of one week. (b) Microfluidic device of 14 μm depth with a growth chamber (6.5 × 2.8 mm^2^) used for monitoring hyphal network growth. Two pillars in the centre prevent the collapse of the growth chamber. 30 microscopic images using a 4× microscope objective was stitched together to monitor chip scale growth.

### Time-lapse microscopy and the Kuramoto model

The spatio-temporal dynamics of the sample was monitored in brightfield using standard microscope objectives with 4x and 10x magnification. The open-source image processing software ImageJ, and custom-written Matlab codes were used to to analyse the captured images [24]. Due to the low height of the microfluidic channel (14 μm depth), hyphal fungal growth in the sample can be indirectly measured by the decrease in image brightness. To analyse the synchronisation of fungal growth dynamics, we estimated fresh growth by subtracting images at 90-minute intervals. The differential brightfield intensity extracted through this methodology was subsequently subjected to detrending procedures, yielding a refined graphical representation of oscillation in spore growth rates. The Kuramoto model was simulated using Matlab and the order parameter was calculated as described in the main text.

## Results

To investigate the spatio-temporal dynamics of fungal growth, *S. indica* cells grown on agar plates for one week (Figure 1a) were inoculated inside a custom-designed microfluidic device (Figure 1b).

### Hyphal growth and network formation inside microfluidic devices

Harvested spores were loaded in a microfluidic device and were supplied with growth media to enable their germination. We sequentially imaged the entire growth chamber to capture images at 30 spatial locations every 10 minutes for over 120 hours. Images were then stitched to obtain macroscale spatio-temporal pictures of hyphal networks (Figure 2(a)). Individual hyphae from separate spores vegetatively fused to create a single large filamentous network. Figure 2 shows representative hyphal filaments from two distinct spores (labelled *S2* and *S3* on the figure) altering their direction midway to align towards a filament from another spore (labelled *S1*), leading to hyphal fusion (Figure 2(b)).

**Figure 2:**
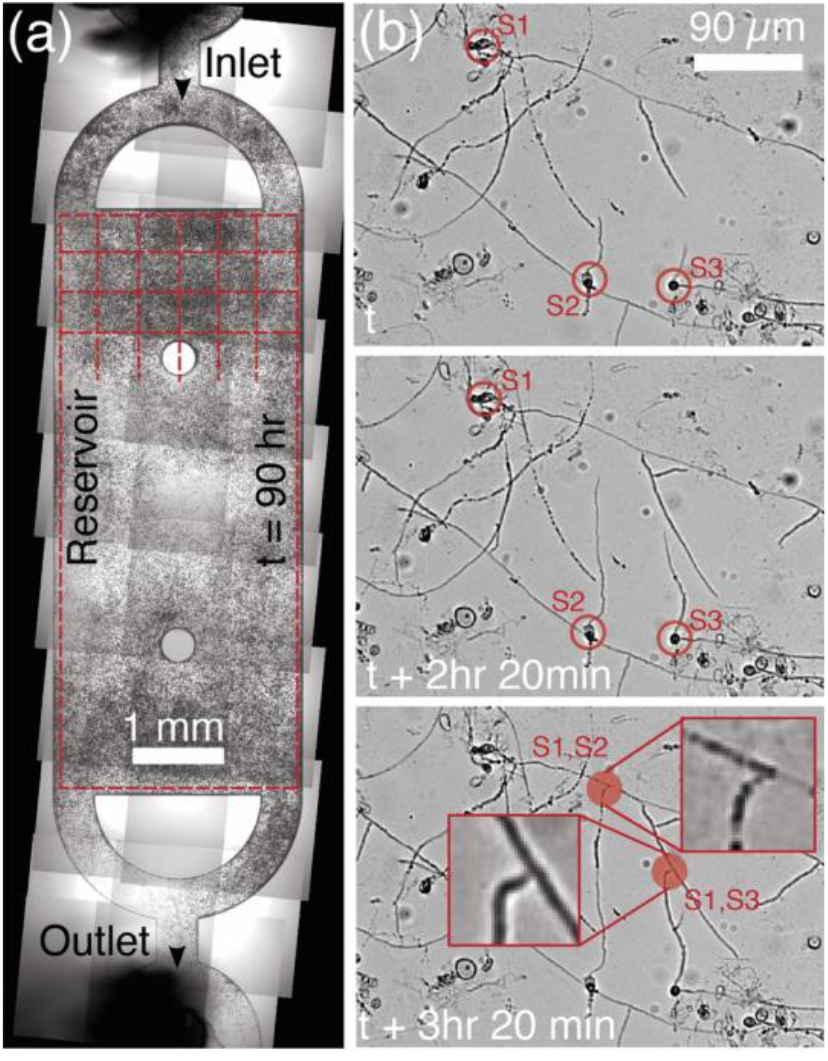
Stitched images of hyphae network in a microfluidic device. (a) Dynamics in the growth chamber is studied by dividing this area in 14 rows and 6 columns of equally sized squares. (b) Hyphae originating from nearby spores fuse to form a filamentous network. Example of hyphae fusion at t + 3hr 20 min.

### Fungal network displays synchronised growth oscillations

After inoculating the microfluidic chamber with *S. indica* spores, it took between 6-24 hrs for spores to germinate, after which the hyphal network spread rapidly (SM Video 1). The diameter of hyphal tips was ∼3 μm and they grew with an intermittent rate of ∼ 0.3-0.4 μm/min. Typically, it took ∼48-72 hrs from the germination of spores to new spore formation (Figure 3(a)). Within 12-14 hrs of first spores appearing, spore size reached a size of ∼10 μm.

**Figure 3:**
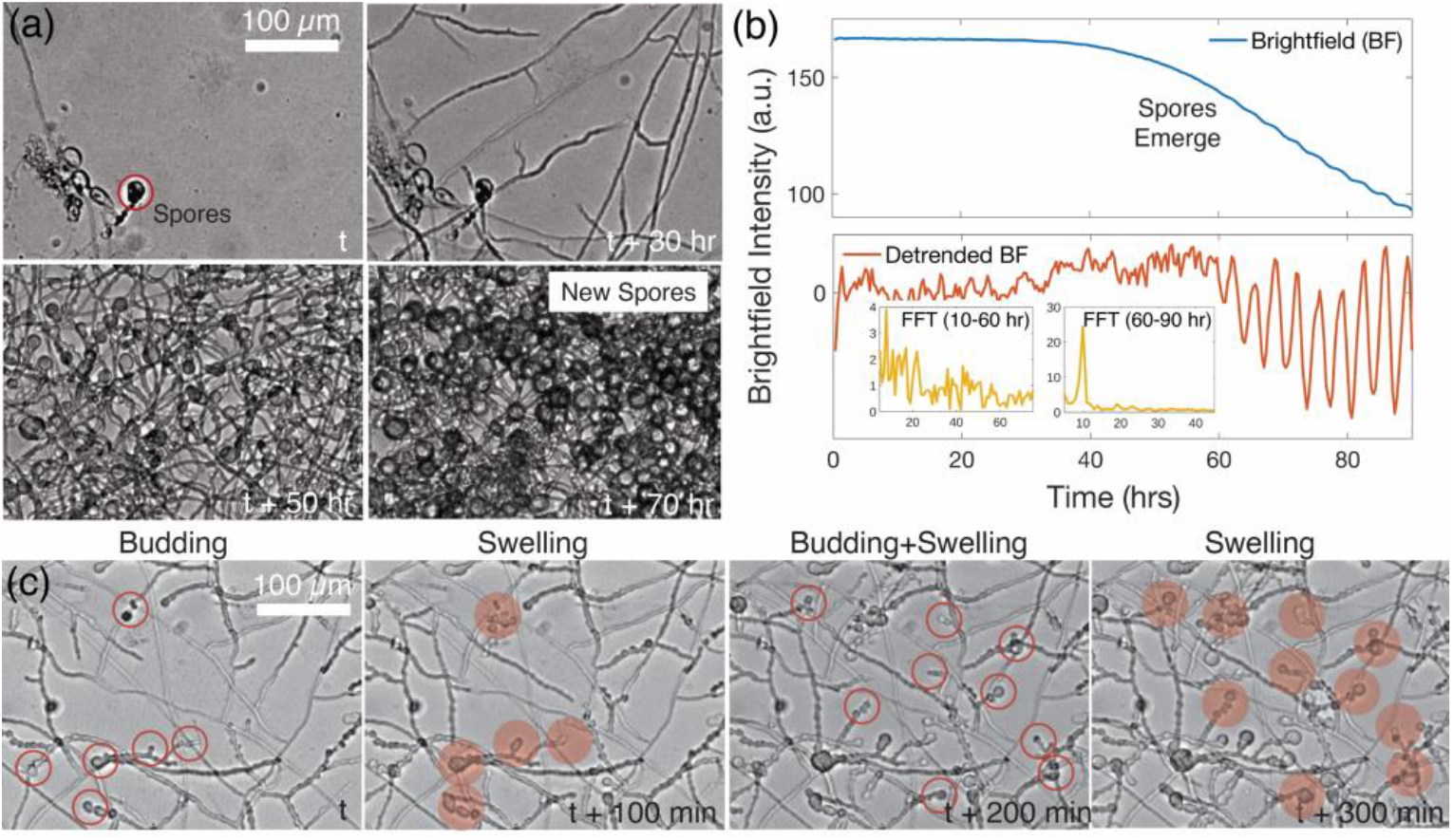
Periodic development of fungal spores. (a) Harvested spores germinating inside a microfluidic device. Over time hyphae grow and form a network followed by the formation of next-generation spores (*timestamp, t* + 70 *hr*). (b) Upper panel: Average brightfield (BF) intensity of hyphal networks shown in (a). Lower panel: Fourier transform of the detrended BF intensity reveals the dominant frequency of oscillations. (c) Next generation of spores periodically form (budding) leading to an increase in the collective growth rate of spores.

The brightfield intensity over time of the growing hyphal network is shown in Figure 3(b). A steep decline in the brightfield intensity after 40 hrs indicates an increased growth rate of fungal biomass. Notably, as the brightfield intensity keeps declining, oscillations with small amplitude appeared after 60 hrs (Figure 3(b)).

Detrending of the brightfield intensity by subtracting average intensity calculated as the arithmetic mean of intensity over two time periods revealed the oscillation patterns (Figure 3(b), lower panel). The frequency of oscillations was calculated from the Fast Fourier transform of the detrended brightfield intensity, which showed a multimodal spectrum until 60 hrs, which ultimately settled to a single frequency corresponding to an oscillation period of 3 hrs (Figure 3(b), inset lower panel). The fungal growth oscillations appeared to be consistent with the periodic budding of spores happening every ∼3hrs as shown in the montage Figure 3(c).

### Synchronization of fungal growth oscillation dynamics

The dominance of a single frequency indicates that the oscillations synchronise across the hyphal networks. To characterise this synchronisation process, we performed a “difference analysis”, where we considered changes in image intensity at 90-minute intervals. The resulting differential brightfield intensity, is expected to be proportional to the fungal growth rate (SM Video 2). To simplify the analysis of the spatio-temporal dynamics, these images were then partitioned into 84 (14 × 6) squares of size 470 × 470 μm^2^. Each region, indicated by its position (i, j) (i=1, ….,14, j = 1, ….,6), was then characterised by the differential intensity signal Δ*I*_*ij*_(*t*) = *I*_*ij*_(*t* + Δ*t*) − *I*_*ij*_(*t*) (Figure 4(a, b)). Following the temporal evolution of these signals reveals the onset of local oscillations at approximately 75 hr after inoculation (Figure 4(c)); average period of ∼3 hr). The amplitude of fungal growth rate is indicated by the intensity gradient of colours from yellow (high) to dark red (low) as shown in the series of images in Figure 4(a). These three images display surface plots of differential brightfield intensity at equally spaced time points. Although, the amplitude varies at individual locations, they remain synchronised as evident from the collective increase in amplitude from minima to maxima. The phase of oscillations at all 84 locations is shown in Figure 4(b) at 75 hrs, when they are scattered, and at 100 hrs, when they are beginning to synchronise.

**Figure 4:**
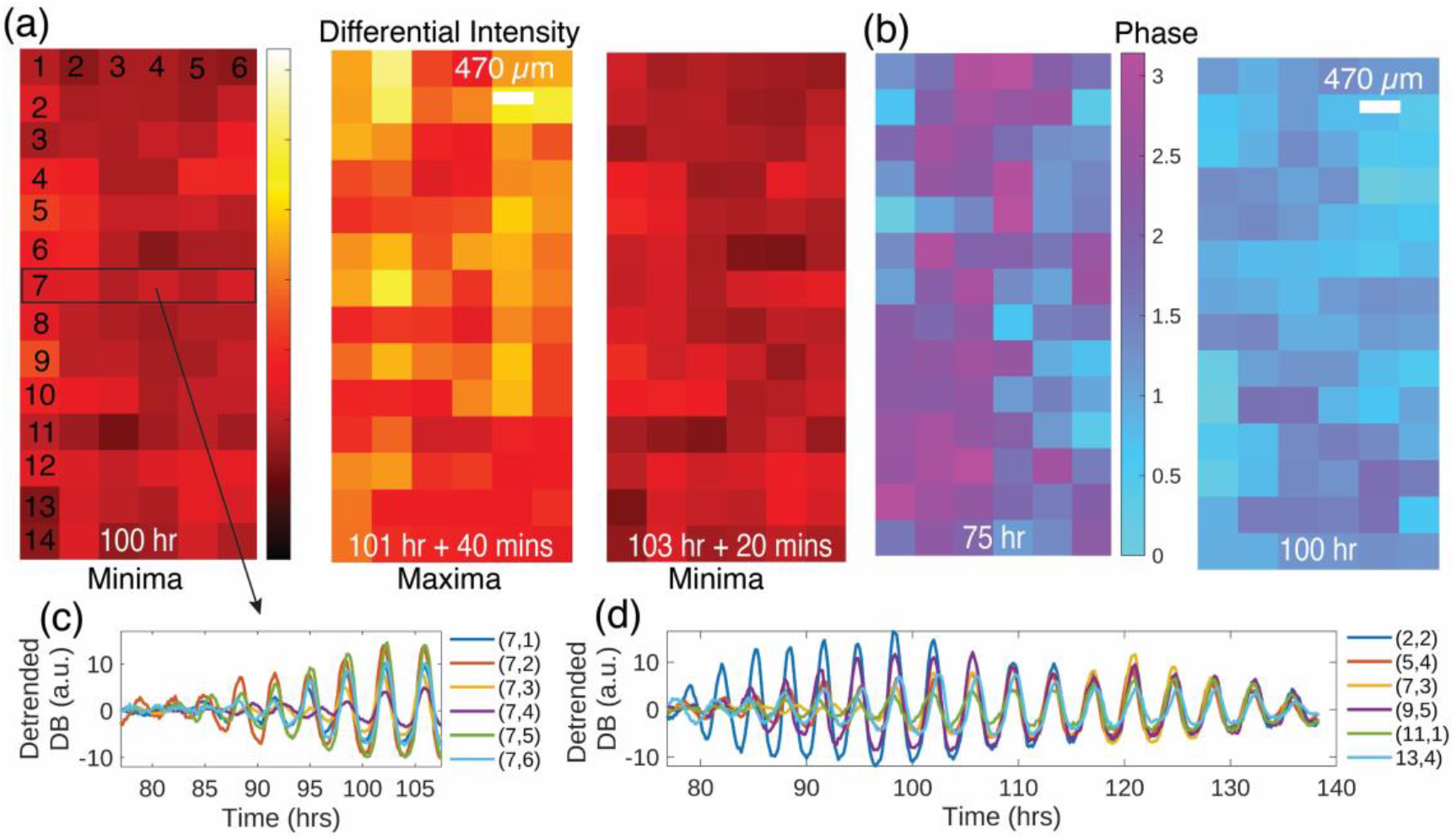
Synchronization of oscillations across a hyphal network. (a) False coloured value of differential brightfield intensity at different locations of the microfluidic device, shown for three different time points as indicated on each panel. The spatial scale is shown on the middle panel. (b) Phase of growth oscillations found in equally sized spatial partitions (14 × 6) across the growth chamber of the microfluidic device. (c) Differential brightfield intensity across six spatial partitions in the 7^th^ row. (d) Differential brightfield intensity at six spatial locations across the growth chamber.

The differential brightfield intensity time series of all the locations in the 7^th^ row {(7, *j*)| *j* = 1, …, 6} and the corresponding detrended, differential brightfield intensity values are shown in Figure 4(c)). Growth rate oscillations at different locations started off with random phases around 75-85 hrs and eventually synchronised around 100 hrs. Growth oscillations at 6 separate locations throughout the growth chamber is shown in Figure 4(d). The oscillations at distinct locations synchronise and remain synchronised for the rest of the observation time. The oscillations appeared on the third day, achieved complete synchrony by the fourth day, and remained synchronised at least until the sixth day (∼140 hrs from inoculation). The global synchronisation of hyphal network throughout the growth chamber suggests that the oscillation dynamics at least up to separation of ∼0.5 cm are coupled.

### Quantification of the synchronisation dynamics

In order to investigate the synchronisation, we begin by estimating the instantaneous phases of the local oscillations *ϕ*_*ij*_(*t*), which were derived from Δ*I*_*ij*_(*t*) whereby the phase of the *n-*th peak at position (*i, j*) is defined as 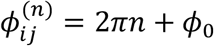 and between successive peaks the phase is taken as linearly increasing. Without loss of generality, we can take *ϕ*_0_ = 0. The frequency of oscillators over the entire microfluidic device - when the growth oscillations are maximally pronounced (102 hrs) - showed the average frequency *ω* = 0.29 (/*hr*) (Figure 5(a)). Experimental phase of the dominant frequency of oscillation at each location was estimated, starting from 75 hrs and for one period (3.5 hrs), then sequentially for later times. Figure 5(b) shows that the distribution of local phases at 75 hr, close to the onset of oscillations, is spread across the whole set of possible values. The phase plot for oscillations began to coalesce close to a single value by 112 hrs at all the 84 locations (Figure 5(c)), signifying a transition to global synchronisation.

**Figure 5:**
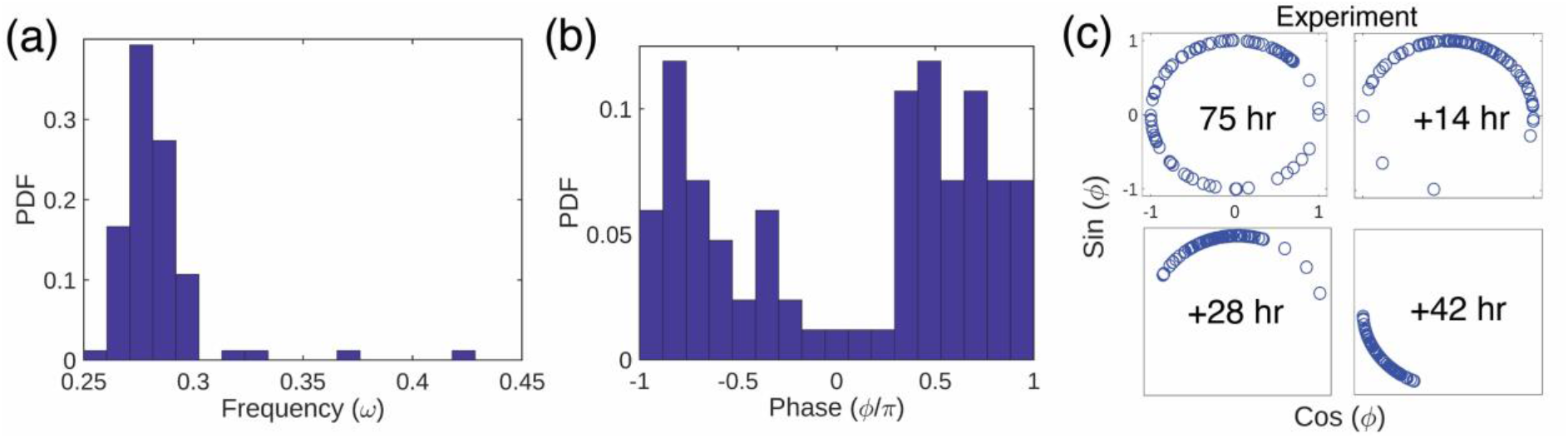
Synchronised oscillation of fungal growth. (a) Distribution of the frequency of oscillators evaluated at 102 hrs, when they are completely synchronised. (b) Distribution of the initial phase of oscillations at 75hrs, when they start to appear. (c) Evolution of the experimental phase of oscillators over time starting from 75 hrs.

### Large-scale synchronisation is recapitulated by the short-range Kuramoto model

To mathematically recapitulate the fungal growth synchronisation dynamics, we used the Kuramoto model, a well-established model known for effectively describing synchronisation in many biological systems [25]. The evolution of the phase of coupled oscillator at the location (a, b) is expressed as:

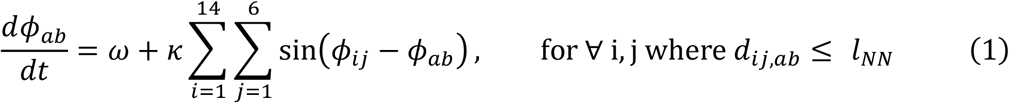

Here, *ω* is the natural frequency, *ϕ*_*ij*_, *ϕ*_*ab*_ are the phase at the location (*i, j*) and (*a, b*) respectively, and *d*_*ij,ab*_ is the Euclidean distance between the location (*i, j*) and (*a, b*). The coupling constant between the oscillators is denoted as κ, and the extent of coupling is invoked when *d*_*ij,ab*_ is smaller than a threshold distance parameter *l*_*NN*_. As the oscillations are coupled it is not possible to estimate the natural frequency of individual oscillator. Therefore, as a first approximation, we treated all oscillators as having the same natural frequency of *ω* = 0.29 which is the average frequency observed at 102 hrs (Figure 5 (a)). Likewise, the initial phase values are estimates at 75 hrs as shown in Figure 5(b). The phase of oscillations is then evolved using nearest neighbour Kuramoto model of the form given in Eq. (1).

To quantify the emergence of synchronisation, the standard order parameter |z|, which measures the phase coherence, was calculated.

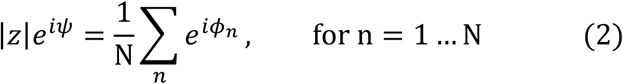

where *ψ* is the average phase of all the oscillators, *ϕ*_*n*_ is the phase of the individual oscillator at a given location, *n* is the individual locations exhibiting oscillations, and N total number of locations. A comparison of evolved phases from the model and the experiments is made by computing the evolution of the order parameter over time using Eq. (2). We run the Kuramoto model simulations with the parameters 0.2 *mm* ≤ *l*_*NN*_ ≤ 1.4 *mm* and 0.025 ≤ κ ≤ 0.9. To compare the simulation and experimental results, the integral area difference between the experimental and simulation order parameter curves over time was calculated for each simulation conditions (Figure 6(a)). In this heatmap, the red region indicates limited synchronisation, and the blue region indicates faster synchronisation compared to the experimental data. The yellow region shows there is a good agreement between experimental and simulation-based evolution of order parameter curves. This parameter region can be approximated by a curve *l*_*NN*_ = 0.3κ^−0.63^, describing the relation between the key parameters of coupling strength and spatial extent (Figure 6(a), dashed line). Figure 6(b) shows the time evolutions of the order parameter in five locations (indicated as A-E in Figure 6(a)). The locations A and C are clearly off from the experimental data as expected from the heatmap. While the locations B and D are along the dashed line, the curves generated with the parameter choices from these regions still showed deviation from the experimental curve (Figure 6(b)). The location E (κ = 0.2 and *l*_*NN*_ = 880 *μm*) most closely recapitulates the experimentally observed order parameter curve (Figure 6(b)). The evolution of the phase with these parameters is shown in Figure 6(c). There appears to be a phase shift when compared to experimental phases shown in Figure 5(c), which can be attributed to the assumptions of constant coupling coefficients and the identical natural frequency for all growth oscillations. Our analysis comparing the experimental and simulation results suggest that the global fungi growth synchronisation across the microfluidic device of an area of 20 mm^2^ are best explained by a model assuming cell to cell coupling within an area of ∼1 mm^2^ [26].

**Figure 6:**
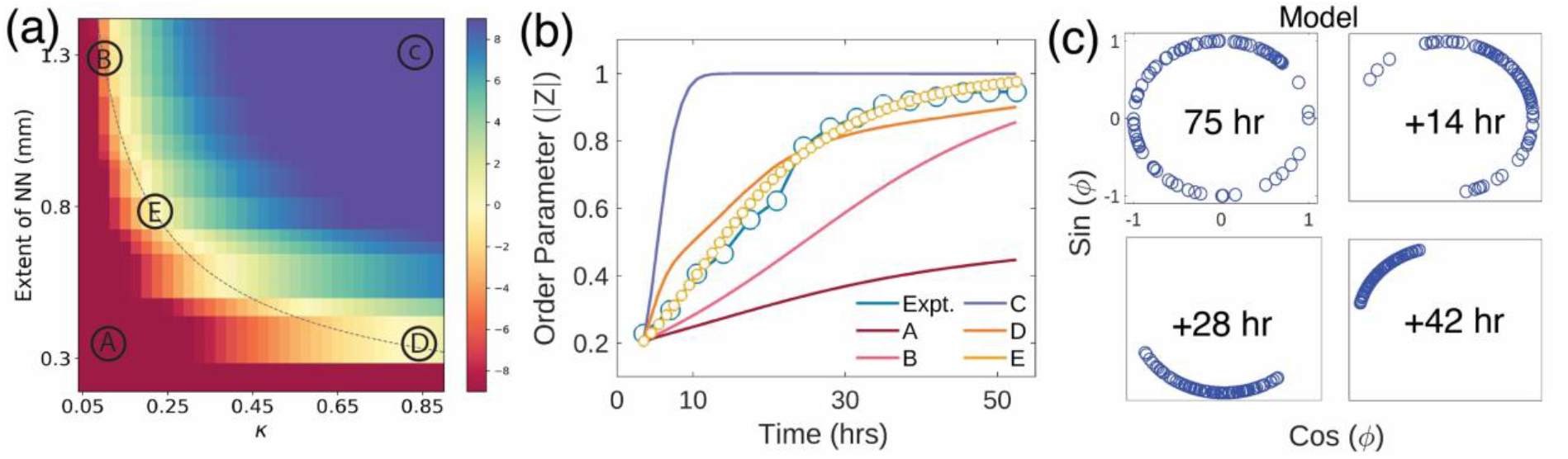
Kuramoto model recapitulates the fungal growth synchronisation. (a) Heat map of the area difference between the experimental order parameter and theoretical order parameter assuming various κ and extent of nearest neighbour coupling *l*_*NN*_. Dashed line is *l*_*NN*_ = 0.3 × κ^−0.63^, which shows a trendline for reasonable agreement with the order parameter and experimental data. (b) Global order parameter from experimental data (Expt.) and the Kuramoto model with the Next-NN coupling with five parameter sets indicated in panel a (A-E). The location E, κ = 0.2 and coupling distance *l*_*NN*_ = 880 *μm*, nearly perfectly retraces the experimental order parameter. (c) Evolution of the phase of oscillators from the model with κ = 0.2 and coupling distance *l*_*NN*_ = 880 *μm*.

## Discussion

We characterized the growth of the plant beneficial filamentous fungus *S. indica* under controlled conditions. By using microfluidic device, we maintained a constant nutrient supply and temperature conditions allowing us to image the growth of fungal colonies for nearly 6 days at a time, indicating that we could monitor the formation of next generation spores that usually occurs 2-3 days after inoculation of harvested spores. We found that fungi display normal growth under these conditions, with the formation of a hyphae network, and that an oscillatory growth pattern emerges after 2.5-days of growth and with the onset of new spore formation. Intriguingly, these oscillations, initially synchronised only locally, result into a global synchronization across a hyphae network covering an area of ∼20 mm^2^. In comparison, the in-plane area of individual hyphae or spores is less than ∼100 μm^2^. Therefore, the total length scale of observed global synchronisation was nearly 200,000 times the size of individual elements responsible for it.

That physio-chemical coupling processes can lead to collective oscillation and traveling waves is well known [29]. For example, collective thickness oscillation in plasmodium is driven by protoplasmic streaming [2], membrane potential oscillations in bacterial biofilm are regulated by spatially propagating waves of potassium [30], and gene expression oscillations in bacteria can be driven by quorum sensing molecules and synthetic transcription networks [31]. Emerging, synchronised oscillations has been previously reported in bacteria *B. subtilis* biofilms, where nutrients are shared periodically between the inside and outside of the colony [27,32]. In that case, the oscillatory dynamics are shown to be mediated by electrical cell-cell signalling [28]. While the underlying biochemical mechanism leading to presented case of synchronisation in *S. indica* hyphae network is unknown, our analysis of the dynamics of synchronisation against a Kuramoto model suggests the existence of coupling at the scale of ∼1 mm^2^. Thus, this works provides the foundation for future studies, to unravel the physiochemical basis of synchronisation in fungal systems.

It remains to be seen if fungal growth oscillations processes are universal to other fungal species. If it is found to be universal, there could be other implications of the collective fungal growth oscillations. In particular, the natural habitat of fungi is soil, which is dense and immobile. Growth waves that propagate through the colony could transport nutrients from the leading front to the interior. As these fungi also colonize plants and form symbiotic relationships with bacteria, it remains to be seen if the collective growth oscillations play a role in the interspecies plant-fungi or bacteria-fungi interactions [33].

## Reference

[1] A. Dussutour, T. Latty, M. Beekman, and S. J. Simpson, Amoeboid Organism Solves Complex Nutritional Challenges, Proc Natl Acad Sci U S A 107, 4607 (2010).

[2] A. Takamatsu, T. Fujii, and I. Endo, Time Delay Effect in a Living Coupled Oscillator System with the Plasmodium of Physarum Polycephalum, Phys Rev Lett 85, 2026 (2000).

[3] T. Nakagaki, H. Yamada, and Á. Tóth, Maze-Solving by an Amoeboid Organism, Nature 407, 470 (2000).

[4] R. Kobayashi, A. Tero, and T. Nakagaki, Mathematical Model for Rhythmic Protoplasmic Movement in the True Slime Mold, J Math Biol 53, 273 (2006).

[5] A. Tero, S. Takagi, T. Saigusa, K. Ito, D. P. Bebber, M. D. Fricker, K. Yumiki, R. Kobayashi, and T. Nakagaki, Rules for Biologically Inspired Adaptive Network Design, Science 327, 439 (2010).

[6] J. D. Julien and K. Alim, Oscillatory Fluid Flow Drives Scaling of Contraction Wave with System Size, Proc Natl Acad Sci U S A 115, 10612 (2018).

[7] M. Kramar and K. Alim, Encoding Memory in Tube Diameter Hierarchy of Living Flow Network, Proc Natl Acad Sci U S A 118, e2007815118 (2021).

[8] M. Tlalka, D. P. Bebber, P. R. Darrah, S. C. Watkinson, and M. D. Fricker, Emergence of Self-Organised Oscillatory Domains in Fungal Mycelia, Fungal Genetics and Biology 44, 1085 (2007).

[9] M. D. Fricker, J. A. Lee, D. P. Bebber, M. Tlalka, J. Hynes, P. R. Darrah, S. C. Watkinson, and L. Boddy, Imaging Complex Nutrient Dynamics in Mycelial Networks, J Microsc 231, 317 (2008).

[10] N. A. R. Gow and M. D. Lenardon, Architecture of the Dynamic Fungal Cell Wall, Nature Reviews Microbiology 21, 248 (2022).

[11] M. L. Smith, J. N. Bruhn, and J. B. Anderson, The Fungus Armillaria Bulbosa Is among the Largest and Oldest Living Organisms, Nature 356, 428 (1992).

[12] J. B. Anderson, J. N. Bruhn, D. Kasimer, H. Wang, N. Rodrigue, and M. L. Smith, Clonal Evolution and Genome Stability in a 2500-Year-Old Fungal Individual, Proceedings of the Royal Society B 285, 1893 (2018).

[13] R. R. Lew, How Does a Hypha Grow? The Biophysics of Pressurized Growth in Fungi, Nature Reviews Microbiology 2011 9:79, 509 (2011).

[14] V. Wernet, M. Kriegler, V. Kumpost, R. Mikut, L. Hilbert, and R. Fischer, Synchronization of Oscillatory Growth Prepares Fungal Hyphae for Fusion, Elife 12, (2023).

[15] A. C. Leeder, J. Palma-Guerrero, and N. L. Glass, The Social Network: Deciphering Fungal Language, Nature Reviews Microbiology 9, 440 (2011).

[16] S. W. Simard, K. J. Beiler, M. A. Bingham, J. R. Deslippe, L. J. Philip, and F. P. Teste, Mycorrhizal Networks: Mechanisms, Ecology and Modelling, Fungal Biol Rev 26, 39 (2012).

[17] M. Held, O. Kaspar, C. Edwards, and D. V. Nicolau, Intracellular Mechanisms of Fungal Space Searching in Microenvironments, Proc Natl Acad Sci U S A 116, 13543 (2019).

[18] L. Heaton, B. Obara, V. Grau, N. Jones, T. Nakagaki, L. Boddy, and M. D. Fricker, Analysis of Fungal Networks, Fungal Biol Rev 26, 12 (2012).

[19] X. Jiang, C. Zerfaß, S. Feng, R. Eichmann, M. Asally, P. Schäfer, and O. S. Soyer, Impact of Spatial Organization on a Novel Auxotrophic Interaction among Soil Microbes, The ISME Journal 12, 1443 (2018).

[20] A. Lareen, F. Burton, and P. Schäfer, Plant Root-Microbe Communication in Shaping Root Microbiomes, Plant Molecular Biology 90, 575 (2016).

[21] A. Fakhro, D. R. Andrade-Linares, S. von Bargen, M. Bandte, C. Büttner, R. Grosch, D. Schwarz, and P. Franken, Impact of Piriformospora Indica on Tomato Growth and on Interaction with Fungal and Viral Pathogens, Mycorrhiza 20, 191 (2010).

[22] X. Qiang, M. Weiss, K. H. Kogel, and P. Schäfer, Piriformospora Indica—a Mutualistic Basidiomycete with an Exceptionally Large Plant Host Range, Mol Plant Pathol 13, 508 (2012).

[23] S. H. Strogatz, From Kuramoto to Crawford: Exploring the Onset of Synchronization in Populations of Coupled Oscillators, Physica D 143, 1 (2000).

[24] J. Schindelin et al., Fiji: An Open-Source Platform for Biological-Image Analysis, Nature Methods 9, 676 (2012).

[25] K. M. Hannay, D. B. Forger, and V. Booth, Macroscopic Models for Networks of Coupled Biological Oscillators, Sci Adv 4, (2018).

[26] M. S. Fischer and N. L. Glass, Communicate and Fuse: How Filamentous Fungi Establish and Maintain an Interconnected Mycelial Network, Front Microbiol 10, 441411 (2019).

[27] J. Liu, R. Martinez-Corral, A. Prindle, D. Y. D. Lee, J. Larkin, M. Gabalda-Sagarra, J. Garcia-Ojalvo, and G. M. Süel, Coupling between Distant Biofilms and Emergence of Nutrient Time-Sharing, Science 356, 638 (2017).

[28] J. M. Benarroch and M. Asally, The Microbiologist’s Guide to Membrane Potential Dynamics, Trends Microbiol 28, 304 (2020).

[29] L. Xiong, A. Garfinkel, L. Xiong, and A. Garfinkel, Are Physiological Oscillations Physiological?, J Physiol 0, 1 (2023).

[30] A. Prindle, J. Liu, M. Asally, S. Ly, J. Garcia-Ojalvo, and G. M. Süel, Ion Channels Enable Electrical Communication in Bacterial Communities, Nature 527, 59 (2015).

[31] A. Prindle, P. Samayoa, I. Razinkov, T. Danino, L. S. Tsimring, and J. Hasty, A Sensing Array of Radically Coupled Genetic ‘Biopixels,’ Nature 481, 39 (2011).

[32] J. Liu, A. Prindle, J. Humphries, M. Gabalda-Sagarra, M. Asally, D. Y. D. Lee, S. Ly, J. Garcia-Ojalvo, and G. M. Süel, Metabolic Co-Dependence Gives Rise to Collective Oscillations within Biofilms, Nature 523, 550 (2015).

[33] P. Bonfante and A. Genre, Mechanisms Underlying Beneficial Plant–Fungus Interactions in Mycorrhizal Symbiosis, Nature Communications 1, 48 (2010).

